# Evolution of flowering time through the asynchrony of pollen dispersal

**DOI:** 10.1101/2023.03.07.531547

**Authors:** Kuangyi Xu

## Abstract

The evolution of flowering time is often attributed to variation of pollinator rates over time. This study proposes that flowering time can evolve through siring success variation among individuals caused by differential pollen dispersal timing (a result of flowering time variation itself). Quantitative genetic models show that earlier flowering evolves under low pollen removal rates, high pollen deposition rates, and a slow decline in the fertilization ability of removed pollen. Variations in pollen dispersal timing also select for a stable variance in flowering time, which is larger when pollen removal rates are either very low or high, pollen deposition rates are moderate, and the fertilization ability of removed pollen declines more rapidly. Also, a model on the coevolution of flower longevity and flowering time predicts that under constant pollination rates, non-random mating results in a weak correlation between late flowering and longer-lived flowers. This baseline finding suggests that the observed correlation between late flowering and shorter flowering duration in nature is influenced by other factors, such as declining pollination rates during late-stage flowering. I discuss the role of altered pollination rates under climate change during flowering time evolution and the importance of distinguishing between pollen removal and deposition rates.

## Introduction

Flowering time is an important trait in plant population since it affects individual reproductive success. Changes in flowering time can be quite rapid in response to environmental changes due to phenotypic plasticity and/or genetic evolution (Simons and Johnston 2000, Fitter and Fitter 2002, Primack et al. 2004, Franks et al. 2007, Büntgen et al. 2022). Flowering time in a population is often continuously distributed, and genetic analyses have found that it has a quantitative genetic basis, controlled by multiple small-effect QTL with moderate to high heritability (Putterill et al. 2004, Buckler et al. 2009, Gonçalves-Vidigal et al. 2009, Xu et al. 2012, Meyer and Purugganan 2013). Since flowering time is a continuous trait, previous works often focused on the evolution of the mean and variance of the flowering time distribution in a population (Elzinga et al. 2007). Models on the evolution of flowering time usually focus on selection caused by temporal variation in pollination rates during the flowering season of a plant population. The temporal distribution of pollination rates is often assumed to be bell-shaped, which may origin from seasonality in pollinator abundance, or that individuals flowering too early or too late may suffer increased competition from other species for generalist pollinators due to displacement of flowering phenologies among species (Mosquin 1971, Heinrich 1975, Rathcke, Elzinga et al. 2007, Devaux and Lande 2009). For the evolution of variance of flowering time in a population, if the reproductive success per individual increases with the density of flowers that are open at the same time, smaller variance may be selected since it increases individual reproductive success by allowing conspecific individuals to flower more synchronously (Fleming 2006). In contrast, larger variance will be selected if the reproductive success per individual decreases with the density of opening flowers (Rathcke 1983, Devaux and Lande 2010). Additionally, flowering time can also be selected by temporal variation in individual viability. For example, in *Silene latifolia*, earlier flowering individuals are more likely to infected by pollinator-transmitted disease (Biere and Antonovics 1996), and in some populations, individuals that flower off-peak will have a lower rate of seed predation (Albrectsen 2000, Mahoro 2002).

The selective factors on flowering time reviewed above rely on temporal variation in external ecological environments. However, flowering time may evolve due to variation in flowering time among individuals itself, which can act even when external environments are constant. Specifically, variation in flowering time may often entail variation in the onset of pollen dispersal and ovule fertilization among individuals. However, under constant pollination rates, variation in the onset of ovule fertilization itself should not cause individuals to have different number of fertilized ovules. In contrast, variation in the onset of pollen dispersal can cause individuals to have different siring success even under constant pollination rates. For example, if removed pollen can persist to sire individuals that flower later, early flowering individuals may obtain higher siring success over late flowering ones by having a long-lasting contribution to the pollen pool, because it can sire ovules that are available both synchronously and later. Indeed, empirical study in *Silene latifolia* found a positive correlation between an earlier onset of flowering and higher male reproductive success (Elzinga 2007). As a result, the mean flowering time of the population may evolve to be earlier. Note that selection on flowering time via variation in the onset of pollen dispersal applies to both hermaphroditic and separate-sex populations, but compared to hermaphrodites, the selective strength will be weaker in separate-sex populations since this selective force acts only through males.

Individual siring success depends on not only its contribution to the pollen pool, but also the number of ovules that are available to be fertilized over time. Moreover, the fertilization ability of removed pollen tends to decline over time, due to the loss caused by pollinator grooming (Devaux et al. 2014) or reduced pollen viability (e.g., reduced vigor, germination, competitiveness with other pollen, Dafni and Firmage 2000). The decline rate of fertilization ability will vary depending on the level of pollen aggregation (Harder and Johnson 2008), and may also differ between annuals and perennials, as well as between wind-pollinated and insect-pollinated species (Culley et al. 2002, Büntgen et al. 2022). As a result, in an outcrossing population, an individual that flowers too early may have low siring success because few ovules are available in the population tencat the beginning of the flowering season, and the fertilization ability of its pollen will have declined to be low when more individuals start flowering and more ovules become available later. Therefore, the evolution of flowering time will be regulated by the distribution of individual contribution to pollen pool and fertilized ovules over time, which should depend on both the distribution of flowering time in the population (e.g., variance of flowering time), and how quickly pollen is removed and deposited, but the effects of these factors on flowering time evolution are unclear.

Variation in the onset of pollen dispersal may also help us better understand the observations that individuals flowering later tend to have shorter-lived flowers (Kehrberger and Holzschuh 2019) or flowering duration (O’Neil et al. 1997, Ollerton and Lack 1998). This correlation may be partly due to variation in individual quality, since later flowering individuals tend to have lower plant and inflorescence height (O’Neil et al. 1997). The correlation may be also a result of temporal variation in pollination rates during a flowering season. However, since pollination rates often drop to be low later in the flowering season, previous models suggest that longer-lived flowers should be favored (Ashman and Schoen 1994), contrary to the observations. This correlation may also be partly generated via non-random mating as a result of variation in the onset of pollen dispersal. Specifically, compared to early flowering individuals, we may expect late flowering individuals to sire fewer ovules since there are fewer ovules available to be fertilized. Since flowers are costly to maintain, individuals with shorter-lived flowers tend to produce more flowers due to the resource allocation trade-off (Ashman and Schoen 1994). As a result, selection may favor late flowering individuals to have shorter-lived flowers with a larger flower number than early flowering individuals. Therefore, to better understand how the correlation among floral traits evolves, it is useful to build a model under the baseline scenario when there is no variation in pollination rates and quality among individuals.

Here I build quantitative genetic models to investigate the evolution of flowering time via differential individual siring success resulted from variation in the onset of pollen dispersal, in the absence of selection caused by temporal variation of pollination rates. Results show that mean flowering time is more likely to evolve to be earlier when the pollen removal rate is lower, the pollen deposition rate is higher, and the fertilization ability of removed pollen declines more slowly over time. Variation in the onset of pollen dispersal among individuals also selects for a certain variation in flowering time that resists the invasion of mutants with either larger or smaller variation. This evolutionarily stable variation of flowering time tends to be larger when the pollen removal rate is either very small or large, the pollen deposition rate is intermediate, and the fertilization ability of removed pollen declines slowly. Based on these findings, I discuss the potential impact of altered pollination rates due to climate change on flowering time evolution, and the importance of distinguishing between pollen removal and deposition rates in empirical studies. Furthermore, under the assumption of constant pollination rates, non-random mating generates a weak correlation between late flowering and extended flower longevity. This suggests that the observed correlation between late flowering and shorter flower longevity in nature may result from other factors, including declining pollination rates in the later stages of flowering and individual quality variations.

## Model

### Model framework

I consider a population with infinitely large population size and non-overlapping generations. For the ease of description, I assume individuals are hermaphroditic, but the model can also apply to separate-sex populations if there is no discordance in the distribution of flowering time between females and males. I assume the population is outcrossing, but the impact of selfing on the results is qualitatively discussed in the discussion section. The model focuses on the evolution of flowering time and flower longevity, and assumes they are continuous traits with a quantitative genetic basis controlled by many small-effect loci, as reviewed in the introduction. The genetic variance of the traits is assumed to be constant over generations. The model assumes no viability selection and does not explicitly incorporate mutations. Flowering time and flower longevity are only selected via differential reproductive success caused by variation in flowering time and flower longevity among individuals, detailed in the following paragraphs. The biological meanings of key symbols used in the model are summarized in Table 1.

**Table 1.** The biological meaning of key symbols used in the model.

| Symbol | Biological meaning |
| --- | --- |
| $t_0$ | Flowering time of an individual |
| $f(t_0)$ | Frequency distribution of flowering time in a population |
| $\bar{t}_0, \sigma$ | Mean and standard deviation of flowering time in a population |
| $R$ | Total reproductive resources for flower construction and maintenance of a plant individual |
| $c$ | Resource cost for constructing a flower |
| $T$ | Flower longevity |
| $m$ | The ratio of the maintenance cost of a flower to the construction cost |
| $F$ | Flower number of an individual |
| $G$ | Female fitness: the proportion of ovules fertilized in a flower |
| $P$ | Male fitness: the proportion of pollen removed in a flower |
| $P_e$ | Effective male fitness after accounting for the fertilization ability of removed pollen |
| $g$ | Female fitness accrual rate: fertilization rate of ovules that remain unfertilized |
| $p$ | Male fitness accrual rate: removal rate of pollen that remain unremoved |
| $r$ | Decline rate of the fertilization ability of removed pollen |
| $w_f$ | Female fitness of an individual |
| $w_m$ | Siring success of an individual, i.e., the proportion of ovules in the population sired by the individual |
| $\tilde{w}$ | Fitness of mutants relative to the residents |
| $u(t_0, T)$ | Joint distribution of flowering time and flower longevity |
| $\bar{T}, \omega$ | Mean and standard deviation of flower longevity |
| $B$ | Genetic covariance between flowering time and flower longevity |
| $V_{t_0, T}$ | Phenotypic covariance between flowering time of the paternal individual and flower longevity of the maternal individual within a zygote |
| $V_{T, t_0}$ | Phenotypic covariance between flower longevity of the paternal individual and flowering time of the maternal individual within a zygote |
| $h_{t_0}^2, h_T^2$ | Heritability of flowering time ($h_{t_0}^2$) and flower longevity ($h_T^2$). |

During a flowering season, I assume an adult individual produces a number of *F* flowers, and all its flowers open simultaneously at time *t*_0_ and fade at *t*_0_ + *T*, during which its pollen is removed and ovules are fertilized. Here *t*_0_ is the flowering time of an individual, which refers to its onset of flowering, and *T* is the flower longevity of its flowers. I assume there is no viability selection on offspring, so selection on flowering time and longevity is solely due to differential fertility among adult individuals. The total fitness of an individual is the production of its flower number and fitness obtained by each flower. The definition of flowering time in the current model is slightly different from previous models. In previous models (Devaux and Lande 2010), flower longevity is not explicitly considered, and flowering time is defined as the time when each flower is open (rather than a trait of an individual). I assume individuals face a resource allocation problem between the flower number and flower longevity (reviewed in Roddy et al. 2021), so individuals with longer flower longevity produce fewer flowers (Ashman and Schoen 1997).

There is variation in flowering time *t*_0_ among individuals, and for most of the results, I assume *t*_0_ is normally distributed as

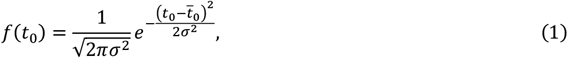

where 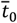 is the mean flowering time and *σ* is the standard deviation. Without loss of generality, it is convenient to set 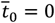. To see whether the results are robust to the distribution function, I also adopted a generalized normal distribution and a Gamma distribution for *f*(*t*_0_) (see section I in Supplementary Materials).

To model the trade-off between flower number and flower longevity, I extended the model of Ashman and Schoen (1994). The model assumes that during one flowering season, individuals have identical limited reproductive resources *R* used for flower production and maintenance, which is assumed to be separate from resources used for fruit and seed provision (Morgan 1992, Schoen and Ashman 1995). However, incorporation of resource allocation between flower and seed/fruit production should not qualitatively change the results, given that it preserves the property that individuals with longer-lived flowers tend to have fewer potential offspring. The resource used for constructing a flower is *c*, and the resource cost for maintaining a flower per time unit is *mc*, where *m* is the ratio of the maintenance cost to the construction cost. Both the construction and maintenance costs are assumed to be invariant over time. Although in some species the maintenance cost may be reduced in the late stage of a flower (Delph and Lively 1989, Weiss 1991), it should not qualitatively change the results as long as the negative association between flower longevity and flower number is preserved. The total resource cost of a flower with longevity *T* is thus *c* + *Tmc*, where *Tmc* is the maintenance cost for a flower with longevity *T*. The flower number of an individual with flower longevity *T* is

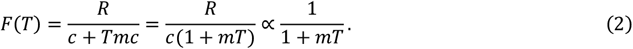

Equation (2) shows that individuals with longer-lived flowers produce fewer flowers due to the resource allocation trade-off. I assume there is no variation in reproductive resources *R* and the cost of flower construction *c* among individuals. Therefore, in the last expression of equation (2), the term *R*/*c* is omitted to reduce the number of parameters (thus it does not mean the number of flowers will be fewer than 1).

The current model follows Ashman and Schoen (1994) to model the pollen removal and deposition process. However, their study focuses on the evolution of flower longevity by assuming individuals bloom at the same time, while the current study focuses on the evolution of flowering time. Following previous literature (Ashman and Schoen 1994, Giblin 2005), I define the female fitness *G* as the proportion of ovules fertilized in a flower, and male fitness *P* as the proportion of pollen removed in a flower. I use the term “siring success” to refer to the proportion of ovules in the population sired by a flower or individual. Note that male fitness is different from siring success because dispersed pollen does not guarantee successful siring of ovules in other individuals. In the model, the siring probability of dispersed pollen depends on the total size of pollen pool and the availability of ovules that receive pollen deposition.

I assume there is no temporal displacement of anther and stigma maturation within a flower (i.e., dichogamy, see Çetinbaş and Ünal (2014) for a recent review), so that pollen dispersal and deposition in a flower starts simultaneously. The effects of dichogamy on the evolution of flowering time can be inferred from the current results and are discussed in the discussion section. For a flower that opens at time *t*_0_, the change of its female and male fitness during an infinitesimal time interval [*t, t* + *dt*] are respectively,

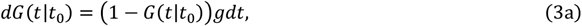

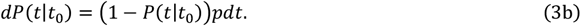

In the above equations, *g* is the fertilization rate of ovules remaining unfertilized until time *t* (i.e., 1 − *G*(*t*|*t*_0_)), and *p* is the removal rate of pollen remaining unremoved until time *t* (i.e., 1 − *P*(*t*|*t*_0_)). *g* and *p* are referred to as female and male fitness accrual rates, respectively (Ashman and Schoen 1994). I assume the fitness accrual rates *g* and *p* are constant over time, so that selection is solely caused by variation in the onset of pollen dispersal. This assumption may not hold in many populations, but it is used to eliminate the confounding selective forces on flowering time caused by temporal variation in pollination rates. The female and male fitness accrual rates may be positively correlated, but may be also discordant in many species (Wilson and Thomson 1991, Young et al. 2007).

By solving equations (3a) and (3b), for a flower that opens at *t*_0_, its female and male fitness change over time as

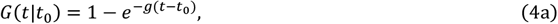

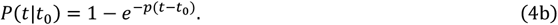

Equations (4a) and (4b) capture the fact that the female and male fitness tend to increase in a decelerating manner over time and saturate at a limit (Wolfe and Barret 1989, Ashman and Schoen 1994). This limit occurs when the maximum potential amount of pollen has been released and when the stigma has been deposited with enough pollen. The calculation of individual siring success is more complicated and is derived in the next subsection.

In the following subsections, I first present models on the evolution of the mean and standard deviation of flowering time by assuming no variation in flower longevity among individuals. I then present the model framework for the coevolution of flowering time and flower longevity when there is variation in flower longevity among individuals.

### Evolution of the mean flowering time

It is useful to first analyze a baseline model for the evolution of flowering time by first assuming no variation in flower longevity. Since in this case, individual will have the same flower number, it is convenient to scale the flower number as *F* = 1. Based on equations (3a) and (3b) and averaging across the distribution of flowering time *f*(*t*_0_), the average proportion of ovules being fertilized and pollen being removed in the population during interval [*t, t* + *dt*] is respectively,

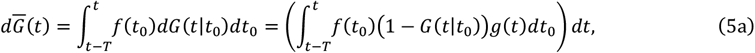

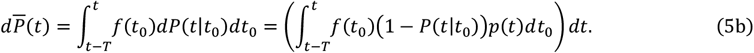

Empirical works suggest that the loss of pollen carried by pollinators (Rademaker et al. 1997, Cresswell 2003, Richards et al. 2009) and the decline of pollen viability (Brunet et al. 2019, Kadri et al. 2022) tend to be decelerating over time. To capture this decelerating decline, I assume the fertilization ability of removed pollen declines exponentially over time as *e*^−*rt*^, where *r* measures how quick the decline is. I refer to the amount of pollen after accounting for the fertilization ability decline as the “effective amount”. The effective amount of pollen in the population at time *t* is the sum over pollen removed before *t* discounted by their fertilization ability, which is

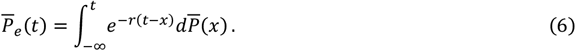

In equation (6), the term 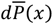 is the proportion of pollen removed during [*x, x* + *dt*], which is discounted by a factor *e*^−*r*(*t*−*x*)^ due to the decline of fertilization ability.

To calculate the fitness of an individual, I use *w*_*m*_ and *w*_*f*_ to denote the fitness an individual obtained through its male and female function, respectively. In other words, *w*_*m*_ and *w*_*f*_ are the number of male and female gametes transmitted to the next generation, respectively. Consider an individual with flowering time *t*_0_ and flower longevity *T*, to calculate its *w*_*m*_, note that its effective amount of removed pollen at time *t* is

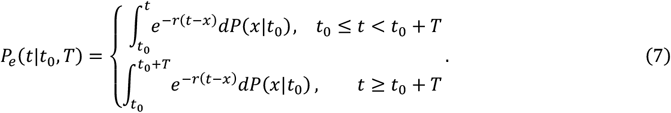

In equation (7), the term *dP*(*x*|*t*_0_) is the proportion of pollen removed during [*x, x* + *dt*], which is discounted by reduced fertilization ability *e*^−*r*(*t*−*x*)^. The upper limit of the integration is *t*_0_ + *T* for *t* ≥ *t*_0_ + *T*, since there is no pollen removed after *t*_0_ + *T* as the flowers have faded. However, the effective amount of pollen contributed by the individual will not be 0 since pollen removed before time *t*_0_ + *T* remain capable of sire ovules. I assume the pollen pool is well-mixed and pollination is random, so for an ovule sired at time *t*, the probability that it is sired by a focal individual is equal to the proportion of its effective pollen *P*_*e*_(*t*|*t*_0_, *T*) out of the total effective amount of pollen 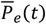(i.e., 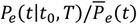). Therefore, the siring success of an individual with flowering time *t*_0_ and flower longevity *T* obtained during time interval [*t, t* + *dt*] is

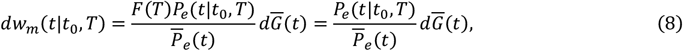

where 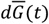 is the proportion of ovules in the population that get fertilized during [*t, t* + *dt*] (given in equation (5a)). The total siring success and female fitness for an individual with flowering time *t*_0_ is 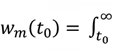 *dw*_*m*_(*t*|*t*_0_*T*) and *w*_*f*_(*t*_0_) = *G*(*t*_0_ + *T*|*t*_0_) (see equation 4(a)), respectively. The overall fitness of the individual, which is the total number of male and female gametes transmitted to the next generation, is *w*(*t*_0_) = *w*_*m*_(*t*_0_) + *w*_*f*_(*t*_0_).

By averaging *w*(*t*_0_) over the distribution of flowering time *f*(*t*_0_), the mean fitness of the population is

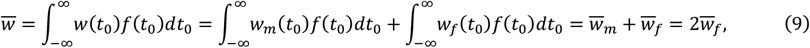

where 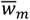 and 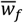 are the mean siring success and female fitness, respectively. The last equality follows from the fact that in a population, the number of pollen and ovules that form offspring zygotes should be the same (thus 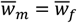). The mean flowering time in the next generation is 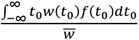, and the selection differential of the mean flowering time (i.e., the difference between the mean flowering time after selection and before selection) is

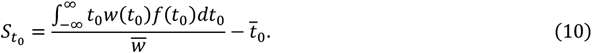

The evolutionary change in mean flowering time after one generation is 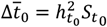, where 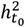 is the heritability of flowering time.

### Evolution of the standard deviation of flowering time

To model the evolution of the standard deviation, I adopt the framework of the evolutionary game theory (Maynard Smith 1982). I consider a residential population invaded by rare mutants. The population size is large enough so that although the frequency of mutants is small, the absolute number of mutant individuals is large. The flowering time distribution of the residents is *f*(*t*_0_), with mean flowering time 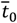 and standard deviation *σ*. For the mutants, the frequency distribution of flowering time is *f*′(*t*_0_) with the same mean flowering time 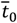 and a different standard deviation *σ*′ among mutant individuals. The mean fitness of the mutants obtained through the female and male functions are respectively

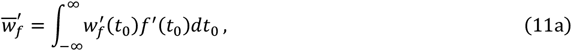

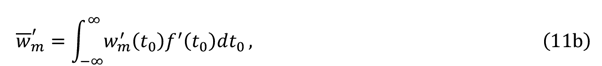

where 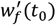 and 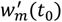 are given in the text below equation (8). Therefore, the relative fitness of the mutants is

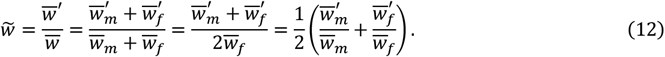

The last two equalities use the fact that the mean fitness of the resident obtained through the female and male functions should be the same (i.e., 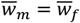, as explained in text below equation (9)). Equation (12) shows that the relative fitness of the mutants is an average of the relative fitness in the male and female function (i.e., 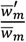 and 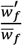). The mutants will invade when 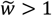. I assume the residents and mutants have the same flower longevity (thus the same flower number), so their mean female fitness will be the same (i.e., 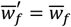). Therefore, the invasibility of the mutants will only depend on whether the mutant on average will sire more ovules than the resident. Assuming that the mutant strategy *σ*′ only deviates from the residential strategy *σ* by a small step, the evolutionarily stable standard deviation of flowering time *σ*\* can be obtained by solving the first-order and second-order conditions and the convergence stability condition (Brännström et al. 2013), which are respectively given by

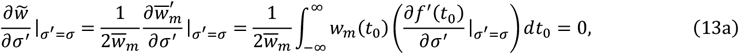

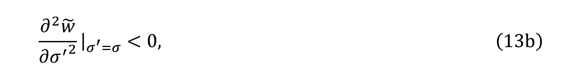

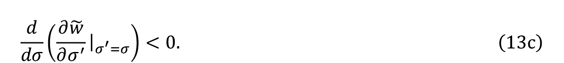

### Coevolution of flowering time and flower longevity

The models in previous subsections assume no variation in flower longevity among individuals, but empirical studies found individuals flowering later tend to have shorter-lived flowers in some species. To understand how this correlation may evolve, I consider the case when there is variation in both flowering time and flower longevity among individuals, but no variation of flowering time and flower longevity among flowers of the same individual. I assume flower longevity *T* is normally distributed with mean 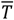 and variance ω^2^. Note that due to resource trade-off, variation in flower longevity will cause individuals to have a different number of flowers open at each time point during a flowering season. Since flower longevity cannot be negative, I assume the distribution is supported on 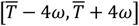 with 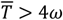, which accounts for 99.9% of a normal distribution. The joint distribution of flowering time and flower longevity is *u*(*t*_0_, *T*).

I use *t*_0,*m*_ and *t*_0,*f*_ to denote the genetic values of flowering time in male and female gametes, and use *T*_*m*_ and *T*_*f*_ to denote the genetic values of flower longevity in male and female gametes. I assume both flowering time and have additive genetic architecture, so that the genetic value of an offspring is the average of parental genetic values. Denote the genetic covariance between flowering time and flower longevity in the parental population by *B*, the genetic covariance in the offspring generation is

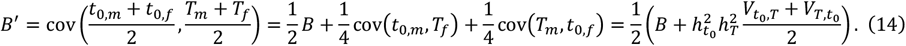

In equation (14), 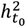 and 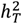 are the heritability of flowering time and flower longevity. 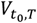is the phenotypic covariance within a zygote between flowering time of the paternal individual and flower longevity of the maternal individual. 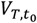 is the phenotypic covariance within a zygote between flower longevity of the paternal individual and flowering time of the maternal individual. Equation (14) implies that the genetic covariance at equilibrium is 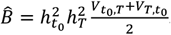. Therefore, whether the correlation between flowering time and flower longevity will negative or positive depend on the sign of the sum of 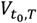 and 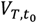.

To calculate 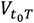, note that during time interval [*t, t* + *dt*], the proportion of ovules from individuals with flower longevity *T* being sired by an individual with flowering time 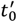 and flower longevity *T*′ is

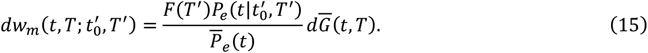

The equation is similar to equation (8). The difference is that in equation (15), for the calculation of the effective amount of pollen in the population 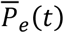 by equation (6), the amount of pollen removed during [*t, t* + *dt*] in the population should be integrated over the join distribution *u*(*t*_0_, *T*) as

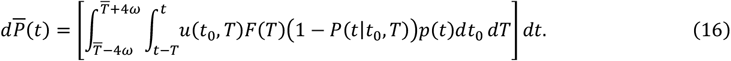

In equation (15), the term 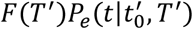 is the effective amount of pollen from the individual at time *t*, where the flower number *F*(*T*′) is given by equation (2), and the effective amount per flower 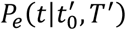 is given by equation (7). 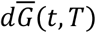 is the proportion of ovules from individuals with flower longevity *T* that get fertilized during [*t, t* + *dt*], given by

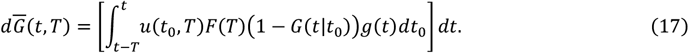

Integrating 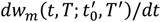 over time *t* and the distribution of flower longevity of the pollen donor *T*′, the proportion of ovules from individuals with flower longevity *T* sired by individuals with flowering time 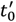 is

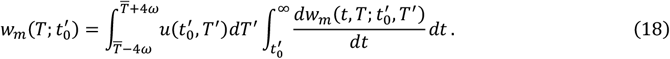

The phenotypic covariance between flowering time of the paternal individual and flower longevity of the maternal individual within a zygote is thus

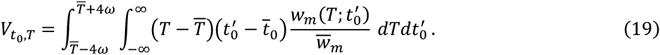

Similarly, to calculate 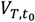, note that during time interval [*t, t* + *dt*], the proportion of ovules from individuals with flowering time *t* being sired by an individual with flowering time 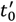 and flower longevity *T*′ is

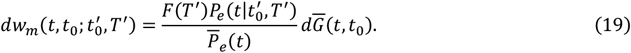

In equation (19), 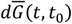 is the proportion of ovules from individuals with flowering time *t*_0_ that get fertilized during [*t, t* + *dt*], given by

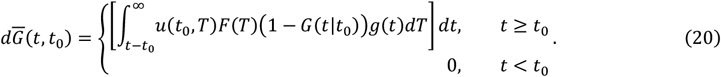

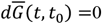 when *t* < *t*_0_ since individuals have not flowered yet. The proportion of ovules from individuals with flowering time *t*_0_ sired by individuals with flower longevity *T*′ is

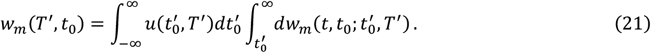

Therefore, the covariance between flower longevity of the paternal individual and flowering time of the maternal individual is

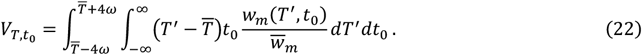

## Results

### Evolution of the mean flowering time

I first investigate the evolution of the mean flowering time by assuming there is no variation in flower longevity. Briefly, the mean flowering time evolves to be earlier when the male fitness accrual rate is low enough and the fertilization ability of removed pollen declines slowly over time (see Fig. 1(a)). Given a certain fertilization ability decline rate of removed pollen *r*, there exists a critical male fitness accrual rate *p* at which the selection differential 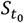 is 0 (illustrated by the bold line in Fig. 1(a)).

**Figure 1.**
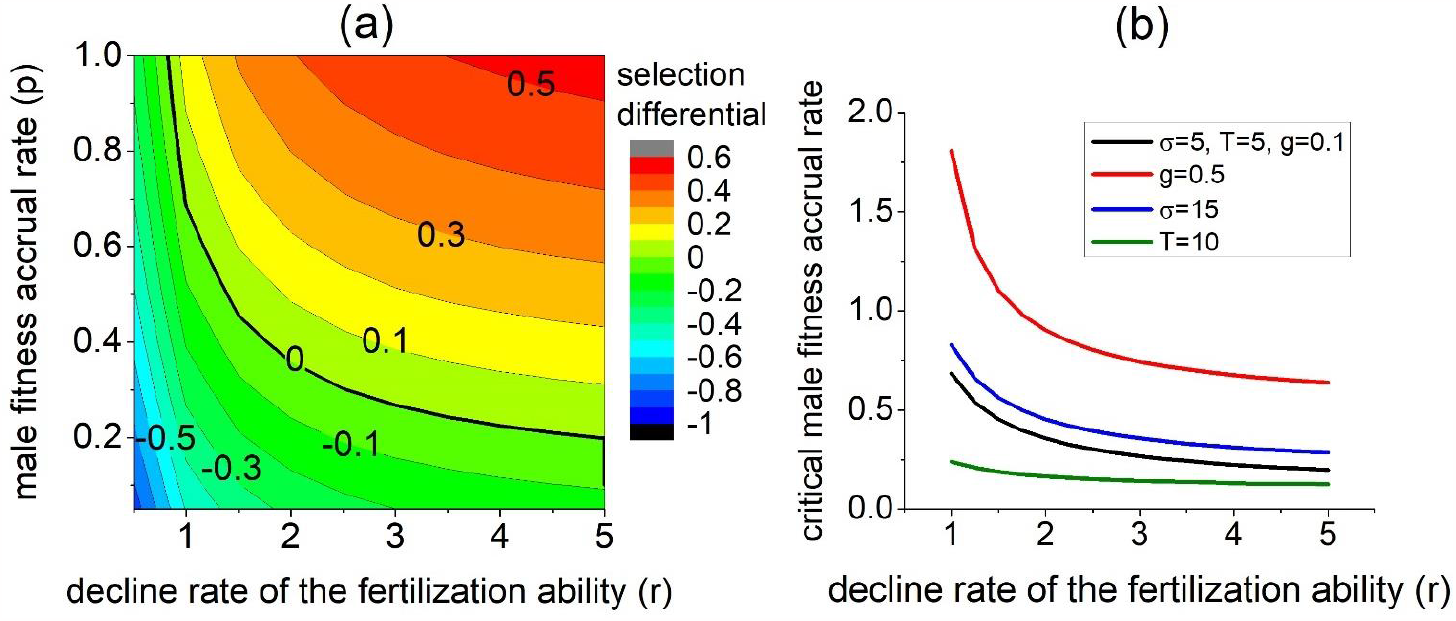
Selection differential of the flowering time 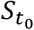 when there is no variation of flower longevity. Panel (a) shows the influences of the decline rate of the fertilization ability of removed pollen (*r*) and the male fitness accrual rate (*p*) on 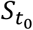, where 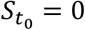along the bold line. Panel (b) shows the effects of several factors on the critical male fitness accrual rate that makes 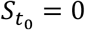, where the black line is taken as the standard condition, and other lines differ from it by one parameter value. Unless specified, *r* = 2, *g* = 0.1, *T* = 5, *σ* = 5.

To understand why a low male fitness accrual rate favors early flowering individuals, I examine the effects of different flowering times on the temporal dynamics of individual contribution to the effective pollen pool 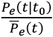 and the rate of change in the siring success 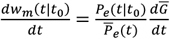 (see equation (8)). Generally, an individual that either flowers early or late will have a higher contribution to the effective pollen pool than an individual with intermediate flowering time (compare the black and blue lines with the red lines in Fig. 2).

**Figure 2.**
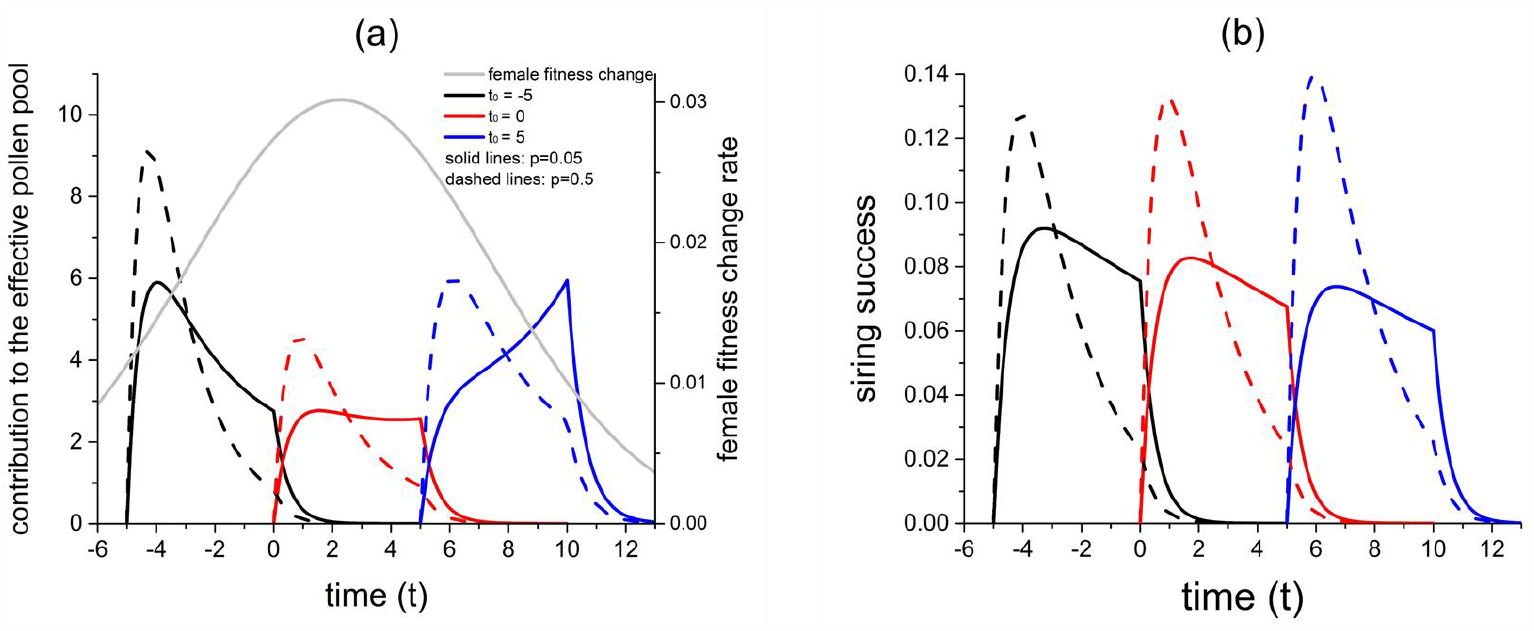
Temporal dynamics of individual contribution to the effective pollen pool 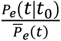 (panel (a)) and the rate of change in siring success 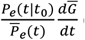(panel (b)). The grey line in panel (a) shows the mean female fitness change rate 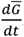. The black, red and blue lines show results for individuals with flowering time *t*_0_= -5, 0, and 5, respectively. The solid and dashed lines show results under low and high male fitness accrual rates, respectively. Parameters are *r* = 2, *g* = 0.1, *T* = 5, *σ* = 5.

However, the siring success also depends on the rate of the female fitness change in the population 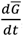, which first increases and then decreases over time (grey line in Fig. 2(a)). Therefore, when the male fitness accrual rate *p* is low, for a late-flowering individual, its contribution to the effective pollen pool peaks at the time when there are only few ovules being fertilized (low 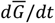, see the blue solid line at *t* = 10 in Fig. 2(a)). As a result, it will obtain relatively low fitness through the male function (compare the three solid lines in Fig. 2(b)). On the other hand, when *p* is high, for an early flowering individual, although its effective pollen contribution rises to a high level at the early stage of its flowering (see the black dashed line at *t* = −4 in Fig. 2(a)), the ovule fertilization rate in the population is low at that time. Moreover, its effective pollen contribution declines more quickly over time compared to later flowering individuals (compare the black dashed at *t* = −2 ∼ 2 with the blue dashed line at *t* = 8 ∼ 12 in Fig. 2(a)). Consequently, an early flowering individual will obtain lower fitness through the male function (compared the three dashed lines in Fig. 1(d)).

Fig. 1(b) shows the influences of other factors on the critical male fitness accrual rate. Mean flowering time is more likely to evolve earlier when the female fitness accrual rate *g* is larger, the standard deviation in flowering time *σ* is larger, and flower longevity *T* is shorter.

To see whether the results are robust to the distribution function of flowering time *f*(*t*_0_), I also considered a generalized normal distribution and a Gamma distribution for *f*(*t*_0_). Generally, the results are robust to different distribution functions (see Fig. S1). Moreover, when *f*(*t*_0_) is a Gamma distribution, which is skewed, the mean flowering time is more likely to evolve to be earlier when the distribution is more right-skewed (Fig. S1(b)).

### Evolution of the standard deviation of flowering time

I investigate the evolution of the standard deviation of flowering time *σ* by assuming no variation in flower longevity. There exists a standard deviation in flowering time *σ*\* that is evolutionarily stable. In other words, mutants with a larger variation (*σ* > *σ*\*) cannot invade a substantial amount of mutant individuals flower earlier or later than the majority of the residents, and thus are unlikely to sire ovules of the residents. On the other hand, mutants with a smaller variation (*σ* < *σ*\*) cannot invade because they are ineffective in siring residential individuals that flower earlier and later than the majority of the mutants. In both cases, the siring success of the mutants will be lower than the residents at the evolutionarily stable strategy.

The invasibility of mutants, thus also the evolutionarily stable standard deviation, depends on how the siring success changes over flowering time. For example, among mutants with larger variation *σ*, compared to the residents, there will be more individuals that flower either very early or late, and thus have either every high or low siring success given that the siring success tends to change monotonically with flowering time (see Figure 2 (b)). Also, there are fewer individuals among mutants that flower around the mean flowering time, which have intermediate siring success. Hence, the invasibility of mutants hinges on whether the overall fitness of an increased population of individuals with either high or low fitness can outweigh a reduced proportion of those with intermediate fitness. This outcome depends on the distribution of individual siring success across flowering time, which is regulated by pollination rates and other factors.

Fig. 3 shows how key factors affect the evolutionarily stable standard deviation *σ*\*. Generally, as the male fitness accrual rate *p* or flower longevity *T* becomes larger, *σ*\* first decreases and then increases, so that *σ*\* will be the smallest at a certain level of *p* or *T* (see Figs. 3(a) and 3(c)). In contrast, as the female accrual rate *g* increases, *σ*\* first increases and then decreases, so that evolutionarily stable variation in flowering time is maximized at an intermediate level of *g* (see Fig. 3(b)). Fig. 3(d) shows that *σ*\* is smaller as the fertilization ability of removed pollen declines more quickly (larger *r*).

**Figure 3.**
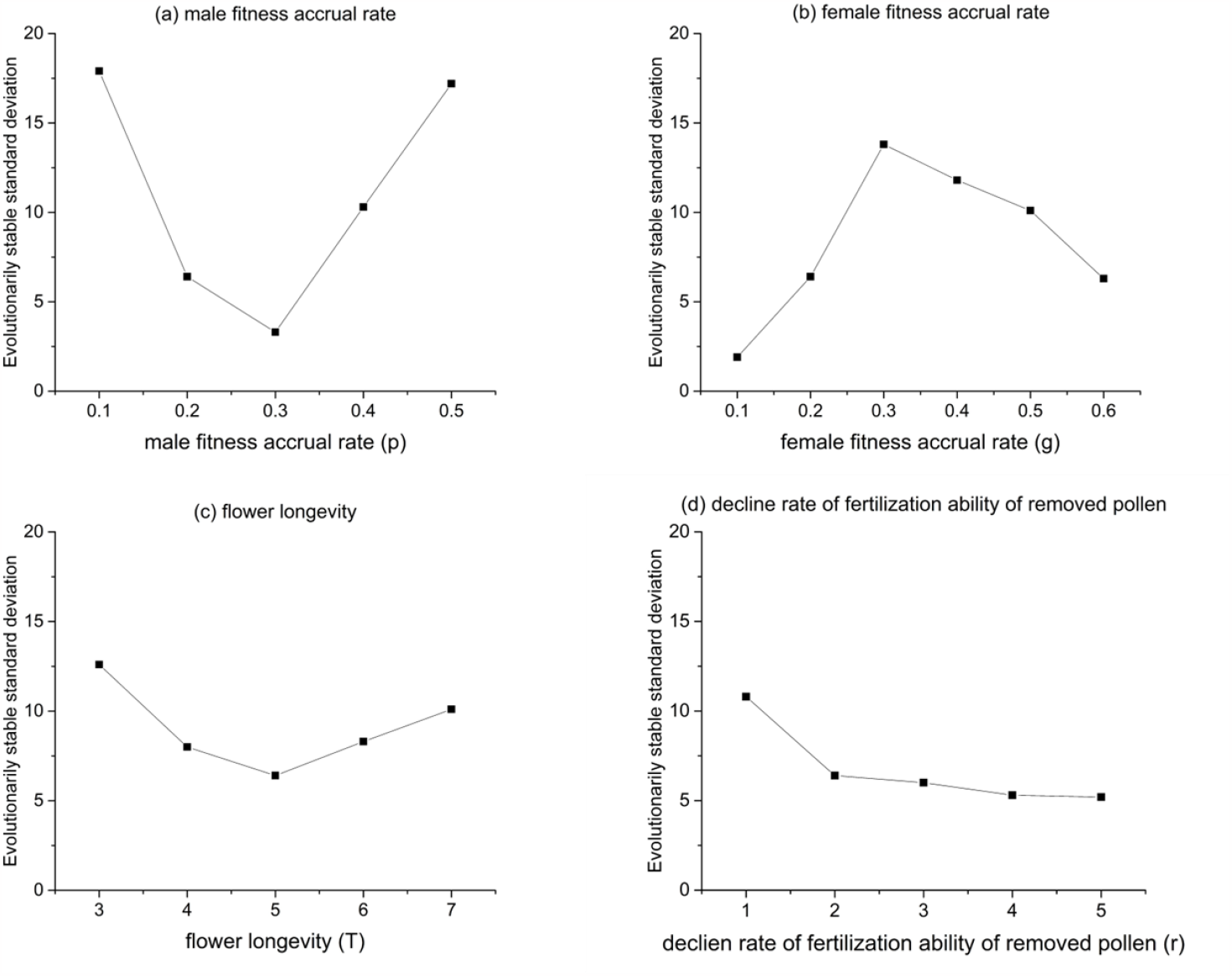
Effects of key factors on the evolutionarily stable standard deviation of flowering time *σ*\* assuming there is no variation in flower longevity among individuals. Unless specified, the parameters used are *T* = 5, *g* = 0.2, *p* = 0.2, *r* = 2.

### Coevolution of flowering time and flower longevity

When there is variation in both flowering time and flower longevity among individuals, equation (14) shows that the equilibrium genetic covariance between the two traits is proportional to the average of two covariances 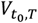 and 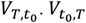 is the covariance between flowering time *t*_0_ of the paternal individual and flower longevity *T* of the maternal individual, and 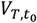 is the phenotypic covariance between *T* of the paternal individual and *t*_0_ of the maternal individual. Under constant pollination rates, the results show that both 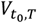 and 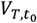 will be positive. Therefore, late flowering individuals tend to produce fewer, but longer-lived flowers.

Biologically, a positive 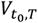 means that early-flowering individuals tend to sire individuals with short-lived flowers. This is because the fertilization ability of removed pollen will decline over time, so during the lifespan of a long-lived flower, it will be mainly sired by individuals that flower later for most of the time. Correspondingly, this correlation is stronger (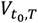 is larger) when the fertilization ability declines more slowly and when the pollen removal is quicker (see Table S1). However, 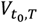 becomes smaller when the female fitness accrual rate is higher, since most ovules in a flower now will be fertilized in the initial stage of its flowering, which causes early flowering individuals to sire more ovules in long-lived flowers.

On the other hand, a positive 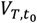 means that individuals with longer-lived flowers tend to sire individuals that flower later. This is mainly because the pollen dispersal process of individuals with long-lived flowers lasts longer. Compared to individuals with short-lived flowers, individuals with long-lived flowers will have a relatively larger contribution to the pollen pool in the later stage of its flowering, because at that time, short-lived flowers have already faded and stop dispersing new pollen into the pollen pool. Due to this cause, 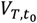 is smaller when the male fitness accrual rate is larger because the male fitness difference between individuals with short-lived and long-lived flowers will become smaller (see Table S1).

An additional investigation on the coevolution of the flowering time and flower longevity (the model is described in section II of the Supplementary Materials) found that variation in flower longevity only very slightly changes the selective strength on the mean flowering time (not shown). Therefore, the results under the assumption of monomorphic flower longevity should qualitatively hold.

## Discussion

This study investigates how the mean and standard variation of flowering time (the onset of flowering for an individual) in a population will evolve due to variation in flowering time among individuals itself. Specifically, variation in flowering time entails variation in the onset of pollen dispersal among individuals, which results in differential siring success among individuals (Elzinga et la. 2007). The study echoes the classical hypothesis that flower production in hermaphroditic flowering plants is primarily controlled by the male function (Sutherland and Delph 1984). In the current study, the ultimate cause of flowering time evolution is the fact that in most plants, pollen (male function) is the transportable reproductive element while ovules (female function) are sedentary, so that the male function of an individual will persist for a longer time than its female function.

Table 2 summarizes the effects of key factors on the evolution of flowering time. Generally, the mean flowering time is more likely to evolve to be earlier when the pollen removal rate is lower, the pollen deposition rate is higher, and variation in flowering time is greater relative to flower longevity. Early flowering is also favored when the fertilization ability of removed pollen declines slowly, as expected. Variation in the onset of pollen dispersal also selects for a standard deviation of flowering time that is evolutionarily stable. At this stable point, mutants with either larger or smaller variation in flowering time cannot invade due to lower siring success than the residents. The size of the evolutionarily stable variation depends on the distribution of individual siring success across flowering time, which is larger when the pollen dispersal rate is either very low or high, the pollen deposition rate is intermediate, or the fertilization ability of removed pollen declines slowly.

**Table 2.** A summary of the effects of key factors on the evolution of the mean and standard deviation of flowering time in a population. The symbols “+” and “−” mean the value of the variable increases and decreases, respectively.

| Flowering time | Male fitness accrual rate ($p$) | Female fitness accrual rate ($g$) | Decline rate of the fertilization ability ($r$) | Flower longevity ($T$) | Variation in flowering time ($\sigma$) |
| --- | --- | --- | --- | --- | --- |
| Mean ($\bar{t}_0$) | $\bar{t}_0 +$ when $p +$ | $\bar{t}_0 -$ when $g +$ | $\bar{t}_0 +$ when $r +$ | $\bar{t}_0 +$ when $T +$ | $\bar{t}_0 -$ when $\sigma +$ |
| Standard deviation ($\sigma$) | $\sigma +$ when $p$ is either small or large | $\sigma -$ when $g$ is either small or large | $\sigma -$ when $r +$ | $\sigma +$ when $T$ is either small or large | N/A |

The results above suggest that changes in pollination rates can have both direct and indirect effects on the evolution flowering time, which can be complicated. For example, an increase in the pollen removal rate directly selects for later flowering time (Fig. 1(a)). However, a high pollen removal rate may select for larger variation of flowering time in the population (Fig. 3(a)) and shorter-lived flowers (Ashman and Schoen 1994), both of which can select for earlier mean flowering time (Fig. 1(b)). Therefore, an increase in the pollen removal rate can have opposing direct and indirect effects on the evolution of mean flowering time, but we may expect the direct effects to act quicker and stronger than the secondary indirect effects.

Climate changes will influence the evolution of flowering time in plants by altering both the level and temporal distribution of pollination abundance, and it is important to understand the relative importance of two selective forces on flowering time: 1) selection by temporal variation in pollinator rates or individual viability (e.g., seed predation, Elzinga et la. 2007), and 2) selection by non-random mating due to variation in the onset of pollen dispersal, as investigated in the current study. For example, under global warming, earlier flowering may be due to earlier pollinator activities, but can also be caused by a decline in the pollen dispersal rate. We may expect the relative strength of the two selective forces on flowering time to depend on the length of flowering periods. Selection by temporal variation in pollination rates may be a dominant force in populations with long flower periods since temporal variation of pollination rates tend to be greater when the flowering period lasts longer. In contrast, populations with short flowering periods may experience more constant pollination rates, so selection caused by variation in the onset of pollen dispersal may dominate.

To see the relative importance of the two selective forces on flowering time under climate changes, one possible way is to compare spring/summer bloomers *versus* fall/winter bloomers. If flowering time is mainly selected by temporal variation in pollination rates, global warming should prompt early flowering in spring/summer bloomers and delayed flowering in fall/winter bloomers. In contrast, if it is changes in pollination abundance that mainly drive flowering time evolution, the two bloom types should show more consistent trends.

It is also important to distinguish between the rate of pollen removal and deposition, since the current results reveal their opposing effects on flowering time evolution (see Table 2). Therefore, how changes in pollination rates will affect the evolution of flowering time should depend on the relative changes in pollen removal and deposition rates. Under pollinator decline, species allocating heavily in the male function may evolve to flower earlier, while those allocating more to the female function may delay flowering, since in the former, pollen removal rate decreases more than the pollen deposition rate. Also, pollen removal and deposition can be performed by different pollinator species (Wilson and Thomson 1991, Young et al. 2007). Therefore, the evolution of flowering time depends on how climate changes affect the relative abundance of these pollinators.

Furthermore, insect-pollinated species may exhibit a slower decline in the fertilization capacity of removed pollen compared to wind-pollinated species (Culley et al. 2002). As a result, when pollination rates decrease, it is more likely for the mean flowering time to evolve earlier, especially in insect-pollinated species. This could partly explain why insect-pollinated species tend to flower earlier than wind-pollinated species in response to climate changes, in addition to the fact that global warming may impact the pollination rates more in insect-pollinated species than wind-pollinated species (Fitter and Fitter 2002).

It is observed that late flowering individuals tend to have shorter flower longevity (Kehrberger and Holzschuh 2019) or flowering duration (O’Neil et al. 1997, Ollerton and Lack 1998). Although the current study shows that under constant pollination rates, non-random mating due to variation in flowering time will generate weak correlation between late flowering and longer-lived flowers, it sets a baseline and suggests that shorter flower longevity in late flowering individuals may result from other factors. One possible cause is pleiotropy due to variation in individual quality, which may be the case in O’Neil et al (1997) and Ollerton and Lack (1998). Specifically, individuals that are less vigorous or in poorer nutritional conditions may grow slower and thus flower later, and these lower-quality individuals also tend to produce shorter-lived flowers. On the other hand, shorter flower longevity in late flowering individuals may also be an adaptive consequence of the declining pollination rate later in a flowering season. This may be the case in Kehrberger and Holzschuh (2019), where there is increased competition and reduced pollination rates over time. In fact, an extension of the previous model (Ashman and Schoen 1994) shows that a temporally declining pollination rate will select for shorter flower longevity, while a temporally increasing pollination rate will favor longer flower longevity (Fig. S2). Moreover, as Devaux and Lande (2010) pointed out, negative correlations observed between onset and duration of individual flowering time (as opposed to flower longevity) may be a statistical artifact caused by variation in flowering duration among individuals.

The current models make several assumptions, some of which can be relaxed by conceptual argument. The models assume an outcrossing population, while self-fertilization is prevalent in plants (Jarne and Charlesworth 1993). Generally, selfing should weaken selection on flowering time but not qualitatively change the results. Depending on the occurrence prior, during or post the cross-pollination process, selfing can be classified into prior, competing, and delayed selfing (Schoen and Lloyd 1992). Under prior or competing selfing, fewer ovules will be available to be cross-fertilized by pollen, which reduces the variation in siring success among individuals. Since delayed selfing occurs after pollination, it will not reduce the ovule availability, which should not affect variation in individual siring success obtained through outcross-pollination. However, self-fertilization of ovules that remain unfertilized after pollination will increase individual fitness (Goodwillie et al. 2018), which reduces the relative fitness differences among individuals. Moreover, selfing may reduce selective responses in flowering time by lowering the genetic variance (Clo et al. 2019). In all, selfing should weaken selection on flowering time, given that the selfing rate does not covary with flowering time.

The difference in selective responses of flowering time between self-incompatible and self-compatible species should be smaller when selection is mediated by temporal variation of pollination rates compared to variation in the onset of pollen dispersal. Specifically, if flowering time is mainly selected by temporal variation in pollination rates, selfing will not affect selective strength on flowering time if selfing relies on pollination (e.g., geitonogamy, de Jong et al. 1993), although selfing may weaken selection if it is autonomous. In contrast, when flowering time evolution is mediated by among-individual variation in the onset of pollen dispersal, all modes of selfing will weaken selection, as discussed before.

The current models also assume no dichogamy (Çetinbaş and Ünal 2014), so pollen dispersal and ovule fertilization start simultaneously in a flower. Dichogamy may affect the evolution of flowering time by causing displacement in the distribution of available ovules relative to the temporal distribution of individual contribution to pollen pool. For example, under protogyny (stigma matures before pollen dispersal), there will be more ovules available at the initial stage of pollen dispersal (i.e., the grey curve in Fig. 2(a) will shift towards left), which should favor earlier flowering individuals by increasing their siring success. Moreover, the time length of dichogamy will evolve by a similar selective forces as flowering time, since variation in the length of dichogamy among individuals can be effectively considered as variation in the onset of the male or female function.

Finally, most studies on the evolution of flowering time focus on the effects of pollination factors. However, meta-analyses indicate that selection caused by abiotic factors may also be important for the evolution of flowering traits (Caruso et al. 2020), and evolution and plastic responses of flowering time to changes in abiotic factors were frequently reported (Franks et al. 2007, Levin 2009, Cho et al. 2017). Therefore, models on selection of flowering time by abiotic factors may be needed. In all, this study shows how flowering time will evolve via non-random mating due to variation in the onset of pollen dispersal among individuals itself. It suggests that climate changes, by altering both the level and temporal distribution of pollinator activities, may have complicated effects on the evolution of flowering time.

## Acknowledgments

This work was supported by the National Science Foundation (Grant No. DEB 1939290).

