## Supplementary Figures and Tables for "Evolution of flowering time through the asynchrony of pollen dispersal"

### Supplementary Tables and Figures for
#### "Evolution of flowering time due to variation in the onset of pollen dispersal among individuals"

**Table S1.** The covariance in the offspring generation with no correlation between flowering time and flower longevity ($B = 0$) in the parental population. $B'/\sigma\omega$ is the change of the correlation coefficient between flowering time and flower longevity after one generation. Unless specified, parameters used are $g = p = 0.2, r = 2, \omega = 1, \sigma = 5, \overline{T} = 10, m = 0.05, h_{t_0} = h_T = 1$.

| | $g = p = 0.2, r = 2$ | $r = 1$ | $r = 5$ | $p = 0.6$ | $g = 0.6$ |
|---|---|---|---|---|---|
| $V_{t_0,T}$ | 0.114 | 0.160 | 0.101 | 0.180 | 0.065 |
| $V_{T,t_0}$ | 0.061 | 0.127 | 0.105 | 0.029 | 0.060 |
| $B'/\sigma\omega$ | 0.008 | 0.014 | 0.010 | 0.010 | 0.006 |

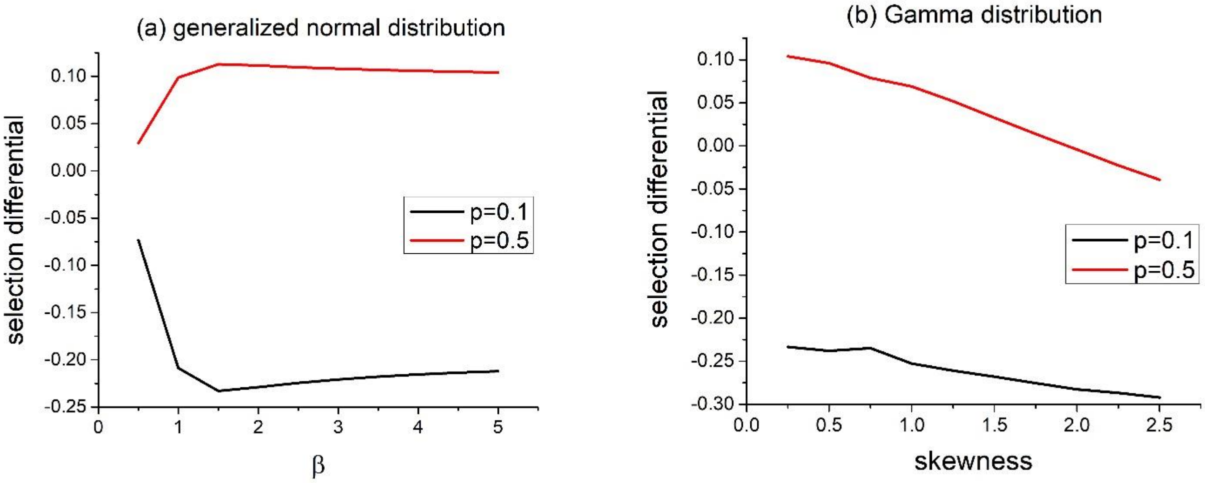


**Figure S1**. Selection differential of flowering time under different distribution functions. Panel (a) show the influences of the parameter $\beta$, which regulates how quickly the frequency decreases with the deviation from the mean. Panel (b) show the effects of the skewness when $f(t_0)$ is a Gamma distribution. The red are results under low and high Parameters used are $r = 2, g = 0.1, T = 5, \sigma = 5$.

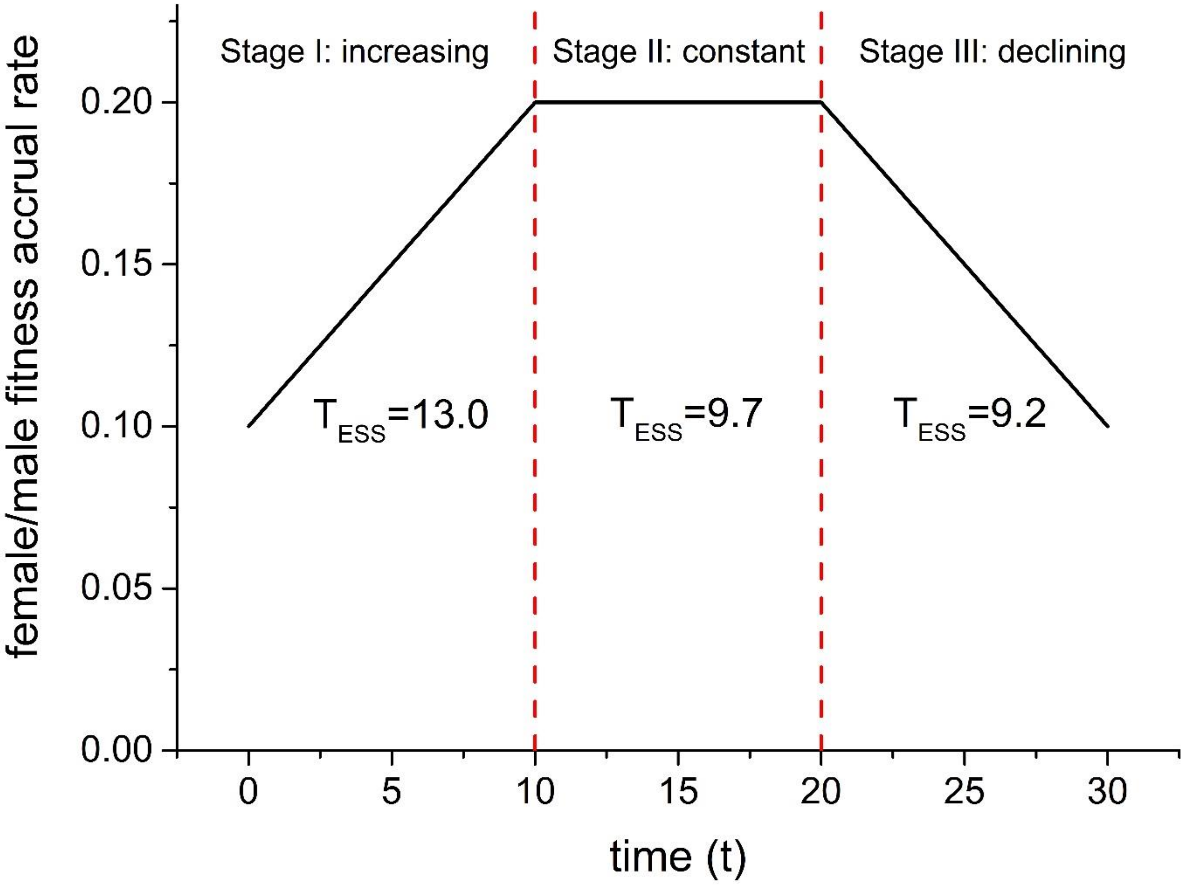


**Figure S2.** The evolutionarily stable flower longevity when the female and male fitness accrual rates change over time. The figure illustrates that during a flowering season, the female and male fitness accrual rates gradually increase (Stage I) and then remain constant (Stage II), and finally decline over time (Stage III). The evolutionarily stable flower longevity $T_{ESS}$ for each stage is shown, which is obtained based on the model of Ashman and Schoen (1994). Although they assume constant fitness actual rates, the model applies to the case of temporally changing fitness accrual rates. The maintenance cost used is $m = 0.05$.
