## Supplementary Materials for "Evolution of flowering time through the asynchrony of pollen dispersal"

**"Evolution of flowering time due to variation in the onset of pollen dispersal among individuals"**

### I. Other distribution functions of flowering time

When $f(t_0)$ is a generalized normal distribution, the expression is

$$f(t_0) = \frac{\beta}{2\sqrt{2}\sigma\Gamma(\beta^{-1})} e^{-\left(\frac{|t_0 - \bar{t}_0|}{\sqrt{2}\sigma}\right)^{\beta}}. \quad \text{(A1)}$$

The mean is $\bar{t}_0$ and the variance is $\frac{2\sigma^2\Gamma(3/\beta)}{\Gamma(1/\beta)}$, where $\Gamma(x)$ is the Gamma function. When $\beta = 2$, $f(t_0)$ is a normal distribution with variance $\sigma^2$. A larger $\beta$ means that the frequency decreases more quickly with the deviation from the mean. To see how the results may change when $f(t_0)$ is asymmetric, I adopt a Gamma distribution, and the skewness of a Gamma distribution with mean $\bar{t}_0$ and standard deviation $\sigma$ is $\mathbb{E}\left[\left(\frac{t_0 - \bar{t}_0}{\sigma}\right)^3\right] = \frac{2\sigma}{\bar{t}_0}$.

### II. Coevolution of the mean flowering time and mean flower longevity

Here I present the model for the coevolution of the mean flowering time and mean flower longevity. By integrating over the join distribution $u(t_0, T)$, the change of the mean female and male fitness during $[t, t + dt]$ in the population is

$$d\bar{G}(t) = \left[\int_{\bar{T}-4\omega}^{\bar{T}+4\omega} \int_{t-T}^{t} u(t_0, T)F(T)\big(1 - G(t|t_0)\big)g(t)dt_0\, dT\right] dt, \quad \text{(A2a)}$$

$$d\bar{P}(t) = \left[\int_{\bar{T}-4\omega}^{\bar{T}+4\omega} \int_{t-T}^{t} u(t_0, T)F(T)\big(1 - P(t|t_0)\big)p(t)dt_0\, dT\right] dt. \quad \text{(A2b)}$$

Consider an individual with flowering time $t_0$ and flower longevity $T$, the number of ovules in the population it sires during time interval $[t, t + dt]$ ($t \geq t_0$) $dw_m(t|t_0, T)$ is given by equation (8). The fitness of this individual obtained from the male function is $w_m(t_0, T) = \int_{t_0}^{\infty} dw_m(t|t_0, T)$. The fitness obtained from the female function is $w_f(t_0, T) = G(t_0 + T | t_0) = 1 - e^{-\int_{t0}^{t_0+T} g(x)dx}$. The individual fitness is thus $w(t_0, T) = w_m(t_0, T) + w_f(t_0, T)$. Averaging over the joint distribution $u(t_0, T)$, the mean fitness of the population is

$$\overline{w} = \int_{\overline{T}-4\omega}^{\overline{T}+4\omega} \int_{-\infty}^{\infty} w(t_0, T) u(t_0, T) dt_0 dT \quad \text{(A3)}$$

The selection differentials of flowering time and flower longevity after one generation are respectively,

$$S_{t_0} = \frac{\int_{\overline{T}-4\omega}^{\overline{T}+4\omega} \int_{-\infty}^{\infty} t_0 w(t_0, T) u(t_0, T) dt_0 dT}{\overline{w}} - \overline{t}_0, \quad \text{(A4a)}$$

$$S_T = \frac{\int_{\overline{T}-4\omega}^{\overline{T}+4\omega} \int_{-\infty}^{\infty} T w(t_0, T) u(t_0, T) dt_0 dT}{\overline{w}} - \overline{T}. \quad \text{(A4b)}$$

Therefore, the evolutionary changes of the mean flowering time and mean flower longevity after one generation are (Lande 1976)

$$\Delta\overline{t}_0 = \frac{G_{t_0}}{\sigma^2} S_{t_0} + \frac{B}{\omega^2} S_T, \quad \text{(A5a)}$$

$$\Delta\overline{T} = \frac{B}{\sigma^2} S_{t_0} + \frac{G_T}{\omega^2} S_T, \quad \text{(A5b)}$$

where $G_{t_0}$ and $G_T$ are the additive genetic variance of flowering time and flower longevity, respectively, and $B$ is the genetic covariance between the two traits.
